# Identification of bacterial signals that modulate enteric sensory neurons to influence behavior in *C. elegans*

**DOI:** 10.1101/2025.09.03.674032

**Authors:** Cassi Estrem, Malvika Dua, Colby P. Fees, Greg J. Hoeprich, Matthew Au, Bruce L. Goode, Lingyi L. Deng, Steven W. Flavell

## Abstract

The bacterial microbiome influences many aspects of animal health and disease. Some bacteria have beneficial functions, for example providing nutrients, whereas others act as pathogens. These bacteria are sensed by host cells to induce adaptive changes in physiology and behavior. While immune and intestinal cells detect bacterial signals through well-characterized mechanisms, recent studies indicate that neurons can also directly sense bacterial signals. However, the bacterial sensory mechanisms in neurons are less well understood. In the nematode *Caenorhabditis elegans*, the enteric sensory neuron NSM innervates the pharyngeal lumen and is directly activated by bacterial food ingestion; in turn, NSM releases serotonin to induce feeding-related behaviors. However, the molecular identities of the bacterial signals that activate NSM are unknown. To identify these signals, we systematically probed bacterial macromolecules from nutritive bacteria using biochemical approaches and GC-MS identification. We find that polysaccharides from gram-positive and gram-negative bacteria are sufficient to activate NSM. We further identify peptidoglycan from gram-positive bacteria as a specific component capable of activating NSM. NSM responses to polysaccharides require the acid-sensing ion channels DEL-3 and DEL-7, which localize to its sensory dendrite in the pharyngeal lumen. Ingestion of bacterial polysaccharides enhances feeding rates and reduces locomotion, matching the known effects of NSM on behavior. We also examine bacterial signals from pathogenic bacteria that can infect and kill *C. elegans*. This approach identifies prodigiosin, a metabolite from pathogenic *Serratia marcescens*, as a bacterial cue that prevents NSM activation by nutritive bacterial signals. This study identifies molecular signals that underlie neuronal recognition of nutritive bacteria in the alimentary canal and competing signals from a pathogenic bacterial strain that mask this form of recognition.

## INTRODUCTION

Animals have coevolved with microbes, leading to complex detection mechanisms that balance immune activation, tolerance of beneficial bacteria, and behavioral adaptations that allow the host to adapt to its microbial environment. The molecular mechanisms by which cells of the immune system and intestine sense bacteria has been the topic of intense investigation over decades. In recent years, it has also become clear that the nervous system plays a direct role in sensing bacteria, though the underlying mechanisms are less well understood.

The molecular mechanisms underlying innate detection of bacteria have been best studied in the immune system. Generic bacterial molecules act as pathogen-associated molecular patterns (PAMPs) that can be detected by pattern recognition receptors (PRRs) on immune cells^1^. Examples of PAMPs include lipopolysaccharide (LPS), peptidoglycan, and microbial nucleic acids; well-studied PRRs include toll-like receptors (TLRs), nucleotide oligomerization domain (NOD)-like receptors (NLRs), and C-type lectin receptors^1–4^. Microbial signals can also be detected by non-immune cells. For example, sentinel cells in the intestinal lining, like enteroendocrine cells, use G protein-coupled receptors to detect bacterial molecules like short-chain fatty acids^5–7^. Recent work has also begun to reveal that specific neuronal cell types can also sense bacterial signals. Nociceptive sensory neurons are directly activated by bacteria during infection, contributing to the perception of pain^8^. Molecular mechanisms in these cells include activation of TRP channels and N-formyl peptide receptors^8–11^. These studies highlight the diversity of molecular mechanisms that different cell types across the body use to sense bacteria. Examining bacterial detection in a wider range of cell types has the potential to identify additional novel mechanisms.

The roundworm *C. elegans* has emerged as a premier model system for the study of host-microbe interactions. *C. elegans* eat diverse bacterial species as their natural diet^12–18^. Bacteria are an appetitive food source to *C. elegans*, providing key nutrients for growth, yet ingestion of pathogenic bacteria can kill the animal^19–25^. As such, *C. elegans* has evolved many mechanisms – exteroceptive and interoceptive -- to sense their bacterial diet and adjust their behavior^26^. Exteroceptive detection involves odorants and tastants from bacteria that can be detected by chemosensory neurons to drive attraction or avoidance to nearby bacterial food sources^27–35^. Interoceptive detection mechanisms are varied. In the enteric nervous system, NSM neurons are activated by the ingestion of appetitive bacteria into the pharyngeal lumen, which triggers feeding, slow locomotion, and persistent dwelling^36–40^. The enteric sensory neuron I3 senses ingested salts to control physiological homeostasis^41^. In the intestine, pathogenic bacteria can damage nearby tissues, which is surveilled by the nervous system to trigger avoidance^23,31,42,43^. Molecules from ingested bacteria can directly affect the host in other ways also. Bacteria-produced neurotransmitters can alter signaling in the *C. elegans* nervous system^44^. Metabolites from bacteria can change metabolic flux in host cells or induce physiological responses that in turn alter neurotransmitter or neuromodulatory signaling to affect behavior^45–47^. Pigments from *P. aeruginosa* bacteria can impact *C. elegans* behavior by changing the spectral content of food lawns^48^. In summary, targeted studies of *C. elegans* cell types with innate bacterial responses have uncovered diverse mechanisms for host-microbe signaling.

NSM is a key hub neuron for interoceptive detection of bacterial food. Previous work showed that ingestion of *E. coli* bacteria directly activates NSM, which in turn releases serotonin to trigger appetitive behaviors, such as feeding and slow locomotion on a bacterial food source^36–40^. NSM detects bacteria via its sensory dendrite that is exposed to the pharyngeal lumen^36,49^. The acid-sensing ion channels (ASICs) DEL-7 and DEL-3 localize to this sensory dendrite and are required for NSM bacterial responses^36^. However, the molecular identity of the bacterial signal that acts on NSM is not known. Given NSM’s central role in inducing feeding-related behaviors, it is possible that it has evolved mechanisms to distinguish nutritive and harmful bacteria, but these possibilities have not yet been explored.

Here, we examine the molecular identities of bacterial signals that act on NSM. We find that NSM responds to a wide range of bacterial species, including gram-positive and gram-negative bacteria. Biochemical approaches reveal that bacterial polysaccharides are sufficient to activate NSM and trigger serotonin-dependent behavioral changes. We identify peptidoglycan from gram-positive bacteria as a specific polysaccharide that activates NSM in a *del-3*- and *del-7*-dependent manner. We also find that a metabolite from a pathogenic bacterium – prodigiosin produced by *S. marcescens* – prevents bacterial activation of NSM and associated feeding behaviors. Our work identifies molecular signals that underlie neuronal recognition of nutritive bacteria in the alimentary canal and competing signals from a pathogenic bacterial strain that mask this form of recognition.

## RESULTS

### Bacterial polysaccharides activate the NSM enteric sensory neuron

We previously found that the ingestion of live *E. coli* strain OP50 by *C. elegans* causes activation of the enteric sensory neuron NSM^36^. This activation is preserved in *unc-13* mutants with defective synaptic transmission but is abolished in mutants with deficits in the outgrowth of NSM’s sensory dendrite^36^. These observations suggest that ingested bacterial food likely activates NSM directly through its sensory dendrite in the pharyngeal lumen. Additionally, we previously found that NSM was not activated by heat-killed bacteria, ingestible beads, or the motor act of pharyngeal pumping, suggesting that its activation is not solely due to mechanical stimulation^36^. As a first step towards identifying the bacterial signal that induces NSM activation, we examined NSM responses to diverse bacterial species, including species commonly used in the laboratory and many others previously isolated from natural *C. elegans* habitats (Fig. 1C)^12^. We examined responses to 12 gram-negative strains and 8 gram-positive strains from diverse phyla (Table S1). To examine NSM responses, we performed *in vivo* calcium imaging of NSM in animals separated by PDMS spacers (Fig. 1A). NSM::GCaMP6m animals were imaged on NGM agar with different bacterial species seeded on the agar surface. We recorded animals during their first exposure to these bacterial species, after a 10-minute acclimation period. To exclude indirect effects of other neurons potentially influencing NSM activity, we performed these experiments in an *unc-13* mutant background, which strongly attenuates synaptic transmission (this is true of all other NSM GCaMP experiments in the study, unless otherwise noted; Fig. 1B shows example traces)^50^. Compared to no-food controls (and LB buffer controls, Fig. S1A), we observed significant increases in NSM calcium in response to all 20 bacterial species tested, though the magnitude of the NSM responses varied (Fig. 1C). Interestingly, different strains of the pathogenic bacterium *Serratia marcescens* activated NSM to different degrees, a result we return to later in the paper. These results suggest that NSM can be activated by ingestion of many different bacterial species.

**Figure 1.**
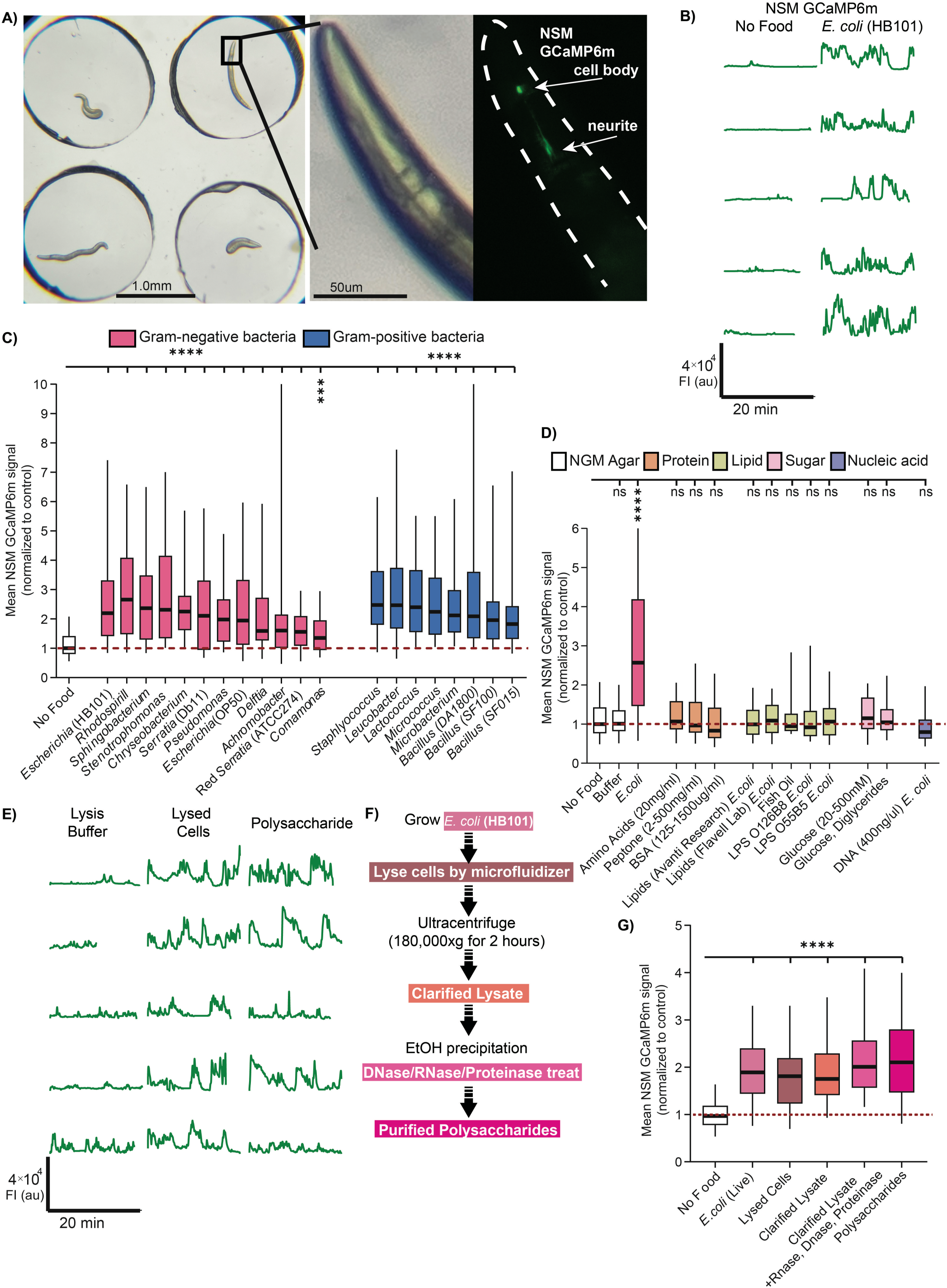
Bacterial polysaccharides activate the enteric sensory neuron NSM. (A) Left: Widefield image of setup for NSM GCaMP imaging on NGM agar with PDMS spacers. Middle: Zoom of *C. elegans* head. Right: GCaMP6m fluorescence in NSM. Note the NSM cell body and the neurites that extend in the posterior direction. All GCaMP quantification throughout the study was performed on the NSM cell body GCaMP signal. (B) Example NSM GCaMP6m traces from *unc-13(s69)* animals in the presence or absence of bacterial food *E. coli* (strain HB101). Each line is a GCaMP recording of one animal (the data are not paired, so different animals are shown for the different experimental conditions). (C) Mean GCaMP signal from NSM in *unc-13(s69)* animals on indicated bacteria. In this plot and subsequent ones, data are normalized to day-matched No Food control recordings. Box plots show median and interquartile range; whiskers show 95 and 5 percentiles. ****p<0.0001; ***p<0.005; ns, not significant Bonferroni-corrected Mann-Whitney test. n = 25-160 animals. (D) Mean GCaMP signal from NSM in *unc-13(s69)* animals on indicated conditions, shown as in Fig. 1C. n.s., not significant by Benjamini-Hochberg FDR-adjusted p-value (q-value) 0.05. ****p<0.0001 by Bonferroni-corrected Mann-Whitney test. n = 15-100 animals. (E) Example NSM GCaMP traces from *unc-13(s69)* animals in the presence or absence of Lysis Buffer, lysed *E. coli* cells or purified Polysaccharide samples. Each line is a recording of one animal. (F) Workflow for bacterial polysaccharide purification. Shades of pink correspond to shades of pink in (G). (G) Mean GCaMP signal from NSM in *unc-13(s69)* animals on indicated conditions, shown as in Fig. 1C. ****p<0.0001 by Bonferroni-corrected Mann-Whitney test. n = 32-83 animals.

Given the broad response profile of NSM, we next tested whether general macromolecules could activate NSM. We examined NSM activity while animals were feeding on NGM agar with different macromolecules on the agar surface: proteins (amino acids, bovine serum albumin), lipids (purified bacterial lipids, LPS, fish oil), carbohydrates (glucose, diglycerides), and bacterial DNA (Fig. 1D; concentrations were determined based on prior work exposing *C. elegans* to these chemicals^51–53^). However, we did not observe activation of NSM under any of these conditions, suggesting that NSM does not generically respond to these macromolecules.

Because NSM’s sensory dendrite is in the pharyngeal lumen, we considered whether these negative results could potentially be due to lower pharyngeal pumping (or feeding) rates during exposure to these macromolecules. If animals were pumping less, they would ingest less of the compound of interest, potentially leading to reduced NSM activity. We tested this by perturbing pumping rates in animals exposed to NSM-activating conditions (live bacteria) or non-NSM-activating conditions (no food, amino acids). To perturb pumping, we used aldicarb, an inhibitor of acetylcholinesterase. Low aldicarb concentrations accelerate pumping by strengthening cholinergic signaling in pharyngeal circuits, while high aldicarb concentrations lead to motor dyscoordination and inhibit pumping^54,55^. Cholinergic signaling is strongly attenuated in *unc-13*, but adding 0.5mM aldicarb still significantly increased pumping (Fig. S1B), and 26mM aldicarb still decreased pumping (Fig. S1C). All animals in our analysis (before and after perturbations) pumped to some degree (ranging from ∼10-150 pumps per minute, in this *unc-13* background). Perturbing pumping rates within this range did not alter NSM activation. For example, live bacteria still activated NSM as effectively when pumping was reduced from 90 to 30 pumps per minute (Fig. S1C, *E. coli* sample). The monosaccharide samples still failed to activate NSM when pumping increased from 18 to 58 pumps per minute (Fig. S1B). We conclude that only a low level of basal pumping is necessary for animals to ingest the contents on the agar surface into their pharyngeal lumen to activate NSM. Consistent with this, we observed that even in no-food conditions (pumping rate of ∼20 pumps per minute), small fluorescent beads on the agar surface were still robustly ingested into the intestine (Fig. S1D,E; consistent with prior work^36^, bead ingestion did not increase NSM activity; Fig. S1F, G). Altogether, these results suggest that minor variation in pumping rates do not impact NSM activity levels and that animals ingest the contents of the agar surface even when pumping at low rates.

Next, we biochemically isolated different components of bacteria and fed them to *C. elegans* during NSM GCaMP imaging. For these studies, we used *E. coli* strain HB101, a commonly used food source for *C. elegans* that evoked one of the strongest responses in NSM (Fig. 1B, C). We found that we could still evoke NSM activity with pressure-lysed HB101 bacteria and a clarified version of this lysate that had been subject to ultracentrifugation (to remove the pelleted debris and dense organelles) (Fig. 1E, G). We next used enzymes to degrade different macromolecules in the clarified lysate and found that treatment with combined DNase, RNase, and Proteinase K did not reduce the magnitude of the NSM response (Fig. 1G; Fig. S2A), suggesting DNA, RNA, and protein are not required for NSM activation. We then switched to a polysaccharide purification method where we ethanol precipitated products from the clarified lysate and treated these products with DNase, RNase, and Proteinase K (Fig. S2A shows an example protein gel illustrating complete protein digestion). Ingestion of the precipitated polysaccharide sample still robustly activated NSM (Fig. 1E-G, Fig. S2B). These results raised the possibility that bacterial polysaccharides are sufficient for NSM activation.

We further examined the precipitated polysaccharide sample to identify its components. Polysaccharides can be hydrolyzed into their monosaccharide building blocks via acid hydrolysis (Fig. 2A). We tested whether this treatment affected NSM activation by subjecting the bacterial polysaccharide sample to acid hydrolysis followed by pH neutralization. Indeed, this attenuated its ability to activate NSM —a result that was also observed when acid hydrolysis was performed on the clarified lysate (Fig. 2B; Fig. S2C). We next subjected the precipitated *E. coli* polysaccharide sample to molecular profiling via gas chromatography-mass spectrometry (GC-MS). To determine the sugar profile of the purified bacterial sample, we subjected two different polysaccharide samples to varying durations of acid hydrolysis and profiled the released monosaccharides. GC-MS analysis confirmed that our purification protocol effectively isolated complex bacterial polysaccharides. As shown in the frequency plot, longer hydrolysis times corresponded to a higher diversity of detected monosaccharides (Fig.2D; Fig. S2E shows another sample with similar results; the samples had different precipitation methods and both activated NSM, Fig. S2B)^56,57^. The released monosaccharides included hexoses (e.g., glucose, galactose), methylated sugars (e.g., rhamnose, fucose), pentoses (e.g., arabinose, xylose, ribose), muramic acid, and additional sugar degradation products such as lactic acid (Fig. 2B-D; Fig. S2E). Based on these results, we tested a more extensive panel of 15 mono- and disaccharides from commercial sources for their ability to activate NSM. This included several sugars that were detected by mass spectrometry, as well as other monosaccharides that are specific to bacteria, such as the main components of cyclic Enterobacterial Common Antigen, Sialic Acid, and lipopolysaccharide (LPS). However, none of them induced NSM activation (Fig. 2E). This further suggests that individual mono- or disaccharides are insufficient to activate NSM. Consistent with this, we separated our purified polysaccharide sample by molecular weight using centrifugal filters and found that the NSM-activating signal in this sample was only retained in fractions larger than 10 kilodaltons (Fig. 2F; monosaccharides are typically <1 kilodalton). Overall, these results suggest that ingestion of large bacterial polysaccharides can induce activation of NSM.

**Figure 2.**
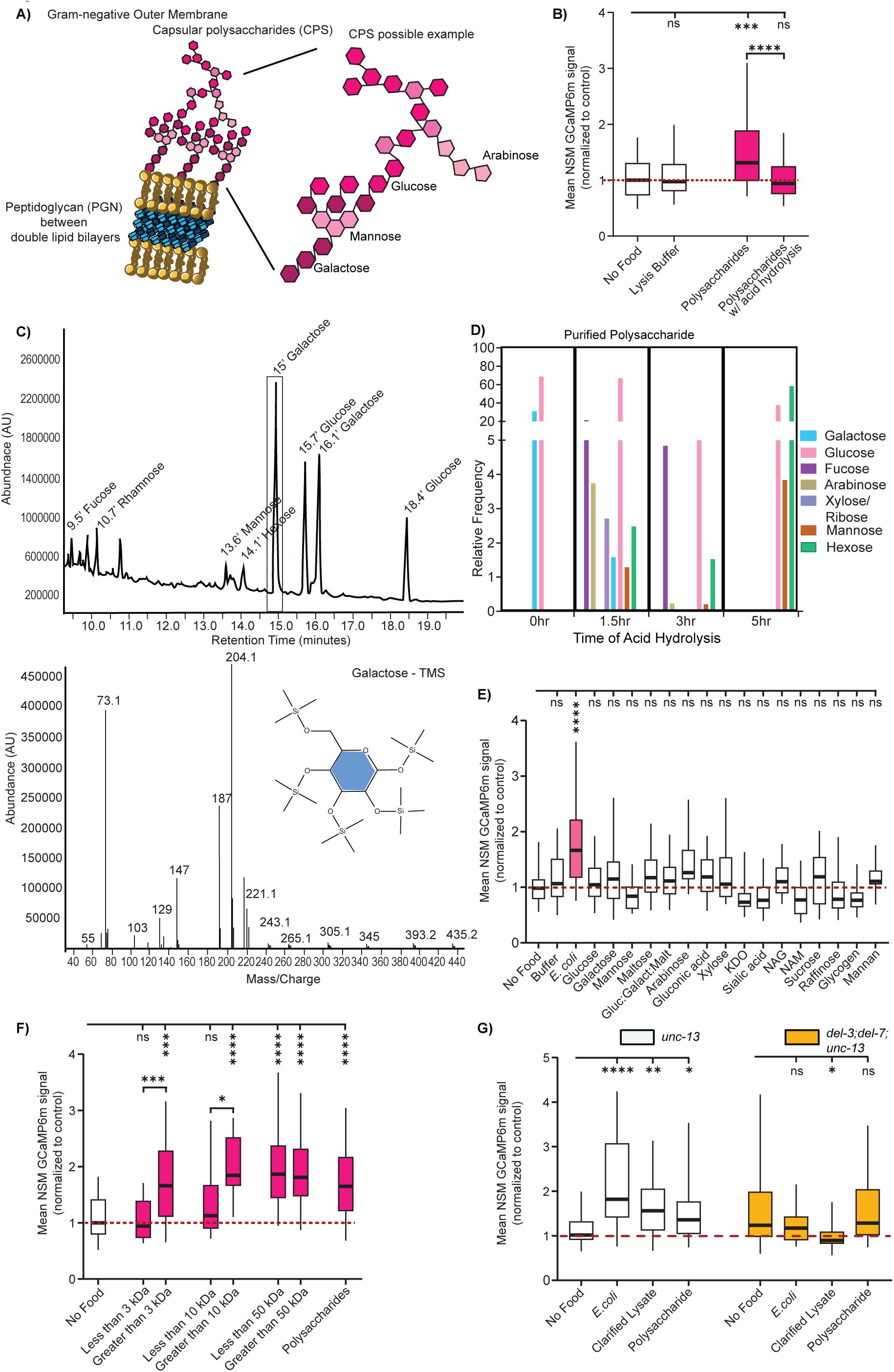
Ingestion of large bacterial polysaccharides activates NSM in a manner that requires ASIC ion channels. (A) Cartoon depicting two sources of polysaccharides from gram-negative bacteria: the outer capsular polysaccharides (monosaccharide components shown in pink) and peptidoglycan layer (in blue) in between the lipid bilayers. (B) Mean GCaMP signal from NSM in *unc-13(s69)* animals on indicated conditions, shown as in Fig. 1C. ****p<0.0001, ***p<0.001 by Bonferroni-corrected Mann-Whitney test. n = 30-137 animals. (C) Top: GC/MS analysis of 70% ethanol precipitated polysaccharide sample from *E. coli* (HB101), subjected to 1.5-hour acid hydrolysis and TMS derivatization. Labels are indicated for eluted monosaccharides identified by GC in which peaks and retention times correspond to those of authenticated standards and MS. Bottom: an example mass spectrum for the major peak at 15’ (galactose-TMS) is shown. Note that different isomers of the same sugar can appear as different peaks. (D) Relative frequency of detected monosaccharides from 70% ethanol precipitation purified polysaccharide from *E. coli* (HB101) hydrolysates at different time points (1.5-hr, 3-hr, and 5-hr) following TFA acid hydrolysis. The y-axis represents the relative abundance (%) of each sugar, normalized to total detected carbohydrates. (a second polysaccharide purification was also subjected to this analysis and is shown in Fig. S2E). (E) Mean GCaMP signal from NSM in *unc-13(s69)* animals, shown as in Fig. 1C. Gluc:Galact:Malt = 30% Glucose: 30% Galactose: 5% Maltose; KDO**=** 3-deoxy-D-manno-octulosonic acid; NAG N-acetyl-D-glucosamine; NAM = N-acetylmuramic acid. ns, not significant by FDR-corrected Mann-Whitney test, Benjamini, Krieger and Yekutieli correction with FDR=0.05. ****p<0.0001 by Bonferroni-corrected Mann-Whitney test n = 13-119 animals. (F) Mean GCaMP signal from NSM in *unc-13(s69)* animals, shown as in Fig. 1C. Samples were passed through centrifugal filters with the indicated size cutoffs. Both the sample that passed through the filter and the sample that was retained were assayed. ****p<0.0001, ***p<0.001 by Bonferroni-corrected Mann-Whitney test. n = 9-172 animals. (G) Mean GCaMP signal from NSM in *unc-13* animals and *del-3;del-7;unc-13*, shown as in Fig. 1C. ****p<0.0001, **p<0.005, *p<0.01; ns, not significant by Bonferroni-corrected Mann-Whitney test. n = 25-121 animals. Note that the significance star in *del-3;del-7;unc-13* is actually a decrease in NSM activity in animals exposed to clarified lysate.

We next tested whether this NSM response to bacterial polysaccharides involves the ASIC ion channels DEL-3 and DEL-7 that localize to its sensory dendrite in the lumen. These channels were previously shown to be required for NSM calcium responses to the ingestion of live *E. coli* bacteria^36^. We tested whether double mutants lacking *del-3* and *del-7* displayed NSM responses to live *E. coli*, clarified bacterial lysate, and the isolated polysaccharide sample. Consistent with prior results, the *del-3; del-7* double mutants failed to display NSM calcium responses to live *E. coli*^36^. In addition, they failed to respond to the clarified lysate and polysaccharide sample (Fig. 2G). This suggests that activation of NSM by ingested bacterial polysaccharides requires the ASIC channels DEL-3 and DEL-7.

### Ingestion of bacterial peptidoglycan activates NSM in an ASIC-dependent manner

We next sought to identify a bacterial polysaccharide of known chemical composition and structure capable of activating NSM. In the above experiments with bacterial polysaccharide isolation, it was challenging to exclude the possibility that a non-polysaccharide contaminant in the sample could be the key activating agent (although certain lines of evidence, for example the acid hydrolysis result, suggested this was not the case). Working with purified monomolecular components would help mitigate this concern. As described above, we had tested lipopolysaccharide from multiple bacterial species, but found that this failed to activate NSM (Fig. 1D). Therefore, we considered other potential candidates.

In addition to LPS, peptidoglycan (PGN) is a prominent bacterial polymer consisting of polysaccharides and amino acids that can signal to host species^58^. PGN is abundant in the cell membranes of gram-positive bacteria. In addition, it is present at lower levels in gram-negative bacteria. It is defined by its repeating NAM-NAG disaccharide units crosslinked to amino acid side chains, though the length of the disaccharide repeats and the nature of the amino acids and crosslinking can differ between bacterial species (Fig. 3A)^58^. We found that feeding *C. elegans* purified PGN from the gram-positive bacteria *M. luteus* or *B. subtilis* led to robust NSM activation (Fig. 3B, C). Feeding *C. elegans* PGN from *E. coli* was not sufficient for NSM activation, suggesting that a different polysaccharide in gram-negative *E. coli* is likely critical for NSM activation (Fig. S2D). PGN from *E. coli* and *B. subtilis* share similar peptide linker lengths and composition; however, *B. subtilis* has a much thicker peptidoglycan layer (10–20 layers compared to 1–3 in *E. coli*), resulting in far more extensive cross-linking^59^. We next tested whether these responses to gram-positive bacteria and their purified PGNs were dependent on the ASICs DEL-3 and DEL-7. Indeed, the NSM response to PGN were absent in *del-3; del-7* double mutants (Fig. 3C). These experiments show that PGN from gram-positive bacteria is a defined polysaccharide that activates NSM in an ASIC-dependent manner.

**Figure 3.**
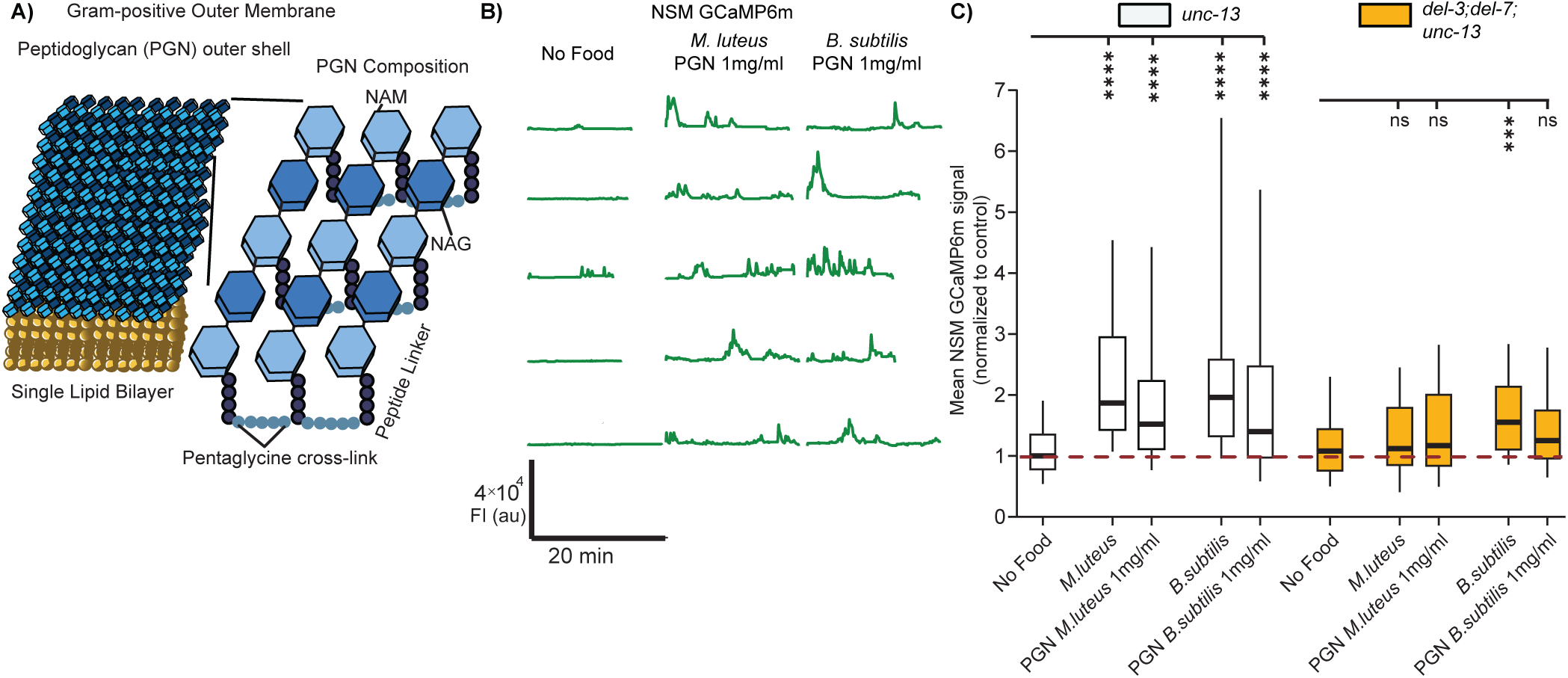
Ingestion of bacterial peptidoglycan activates NSM in an ASIC-dependent manner. (A) Cartoon depicting outside peptidoglycan (PGN) layer in gram-positive bacteria. PGN is made up of repeating NAM-NAG disaccharides with a peptide linker that varies depending on the bacterial species. (B) Example NSM GCaMP traces from *unc-13(s69)* animals in the presence of purified PGN from *M. luteus* and *B. subtilis* (from Millipore Sigma). Each line is a recording of one animal (the data are not paired, so different animals are shown for the different experimental conditions). (C) Mean GCaMP signal from NSM in *unc-13* and *del-3;del-7*;*unc-13* animals, shown as in Fig. 1C. ****p<0.0001; ns, not significant by Bonferroni-corrected Mann-Whitney test. n = 19-123 animals.

### Bacterial polysaccharides and PGN induce behavioral changes associated with NSM activation

NSM activity is associated with behavioral changes typically observed in feeding animals^36–39^. Specifically, optogenetic NSM activation drives increased pharyngeal pumping (or feeding) and slow locomotion^36,60^. NSM inhibition or loss of NSM’s neurotransmitter serotonin (via mutation of *tph-1*, a rate-limiting enzyme for serotonin biosynthesis) partially impairs the increased feeding and decreased speed that animals display on *E. coli* food^36,37^. We examined the effects of *E. coli* polysaccharides and peptidoglycan on behavior*. C. elegans* animals were grown *on E. coli* OP50, then transferred to NGM agar with the indicated components, and behavior was quantified within one hour. Interestingly, we observed elevated feeding rates and reduced locomotion when animals were exposed to either *E. coli* polysaccharides or PGN from the gram-positive bacteria *M. luteus* or *B. subtilis* (Fig. 4A,C; control experiments ruled out the possibility that these effects were due to higher osmolarity of polysaccharide sample, Fig. S3A,B; *M. luteus* and *B. subtilis* were tested because their purified peptidoglycan was commercially available). The magnitudes of these effects nearly matched those observed on live bacteria. We next tested whether the polysaccharide- and peptidoglycan-induced changes in feeding and locomotion required serotonin, the key neurotransmitter in NSM^37^. Indeed, the elevation in pumping and reduction in locomotion were both attenuated in *tph-1* mutants (Fig. 4B, D). These results suggest that *E. coli* polysaccharides and PGN from gram-positive bacteria induce a serotonin-dependent change in feeding and locomotion.

**Figure 4.**
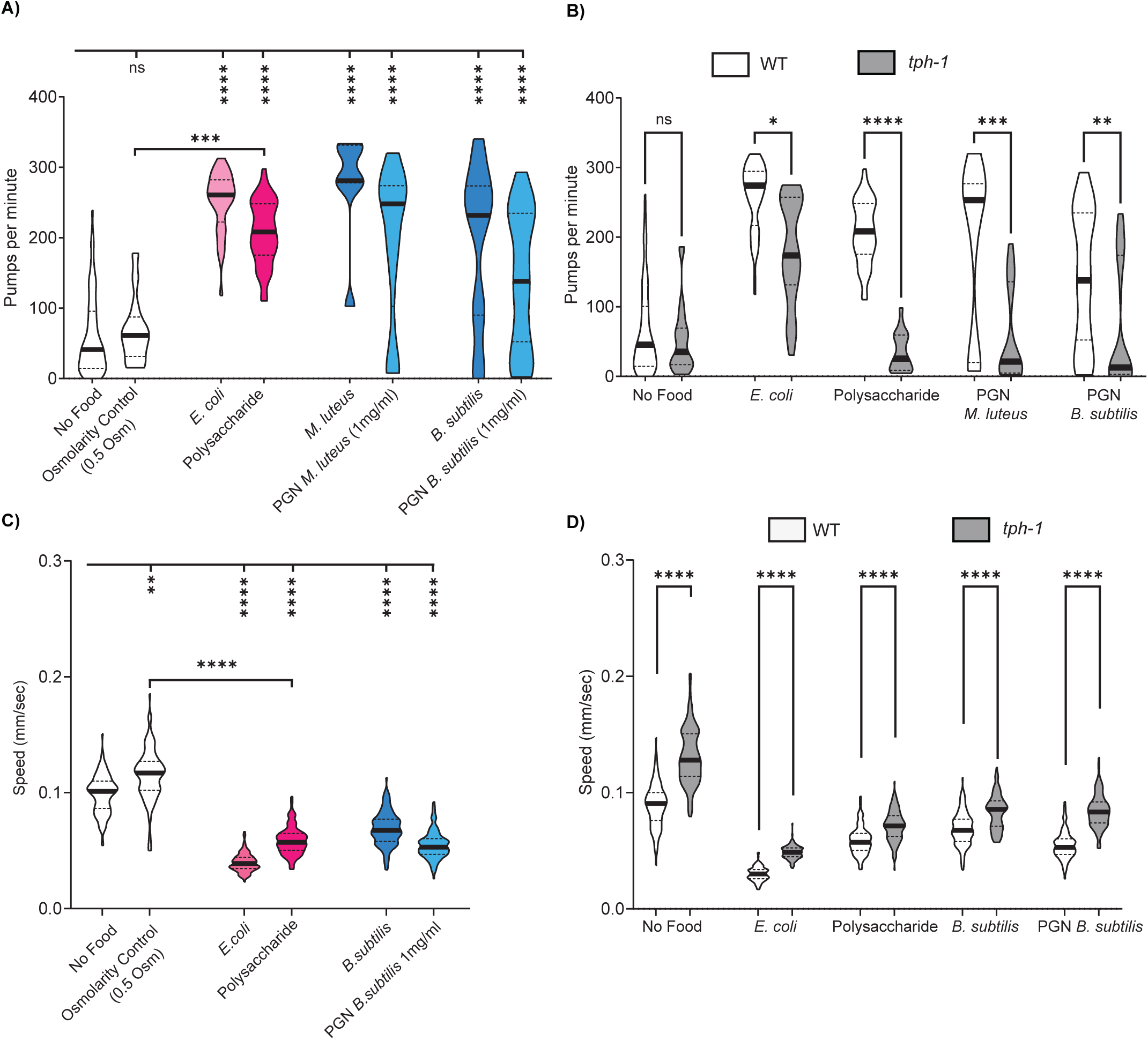
Bacterial polysaccharides, including peptidoglycan, induce behavioral changes associated with NSM activation. (A) Pharyngeal pumping rates of WT (N2) animals exposed to the indicated substances. Violin plot shows median with the solid black line and quartiles with black dashed line. ****p<0.0001, ***p<0.001, ns, not significant by Bonferroni-corrected Mann-Whitney test. n = 25-85 animals. (B) Pharyngeal pumping rates of WT (white) and *tph-1* (grey) animals exposed to the indicated substances for one hour, shown as in Fig. 4A. Note that the WT pumping rates here are the same data as in (A). ****p<0.0001, ***p<0.001, **p<0.005; ns, not significant by Bonferroni-corrected Mann-Whitney test. n = 20-84 animals. (C) Mean speed of WT animals on indicated substances, assayed 15-75 minutes after transfer from growth plates. The violin plot displays the median (solid black line) and quartiles (dashed black lines). ****p<0.0001, ***p<0.001, **p<0.005, *p<0.01; ns, not significant by Bonferroni-corrected Mann-Whitney test. n = 35-125 animals per condition across at least three independent days. (D) Mean speed of WT (white) and *tph-1* (gray) animals exposed to the indicated substances, shown as in Fig. 4C. Note that the WT speed shown here is the same data as in (C). Significance levels: ****p<0.0001, ***p<0.001, **p<0.005, *p<0.01; ns, not significant by Mann-Whitney test. n = 35-125 animals per condition across at least three independent days.

### A *Serratia marcescens* metabolite, prodigiosin, inhibits NSM activity and associated behaviors

We next turned our attention to the microbial cues produced by pathogenic bacteria. As described above, in our initial screen of how different bacteria activate NSM, we noted that different strains of the pathogenic bacterium *S. marcescens* had different effects on NSM. Specifically, *S. marcescens* strain Db11 activated NSM, but *S. marcescens* strain ATCC274 was far less effective (Fig. 1C). These strains are notably different in their appearance. *S. marcescens* (Db11) is non-pigmented (Fig. 5A), but *S. marcescens* (ATCC274) grown at 23C produces a red pigment called prodigiosin (Fig. 5B, C). Previous studies have noted a general trend where red-pigmented *Serratia* tend to be more effective at killing *C. elegans* than non-pigmented strains^61^, but both non-pigmented and red-pigmented *Serratia* can be infectious. The pigmentation of *S. marcescens* strain ATCC274 can be controlled by the ambient temperature during its growth: growing it at 37C leads to non-pigmented bacteria (‘non-pigmented Serratia’), whereas growth at 23C leads to red pigmentation (“Red Serratia”) (Fig.5A-C)^65,66^. Prompted by our initial results from the screen, we compared how well *S. marcescens* ATCC274 grown at these two temperatures activates NSM (Fig. 5D; the actual GCaMP imaging was performed at room temperature, shortly after bacterial growth). This revealed that the non-pigmented *Serratia* robustly activated NSM, whereas the Red *Serratia* did not. These results suggest that this single bacterial strain can have different effects on NSM activation depending on its growth and, thus, its chemical makeup.

**Figure 5.**
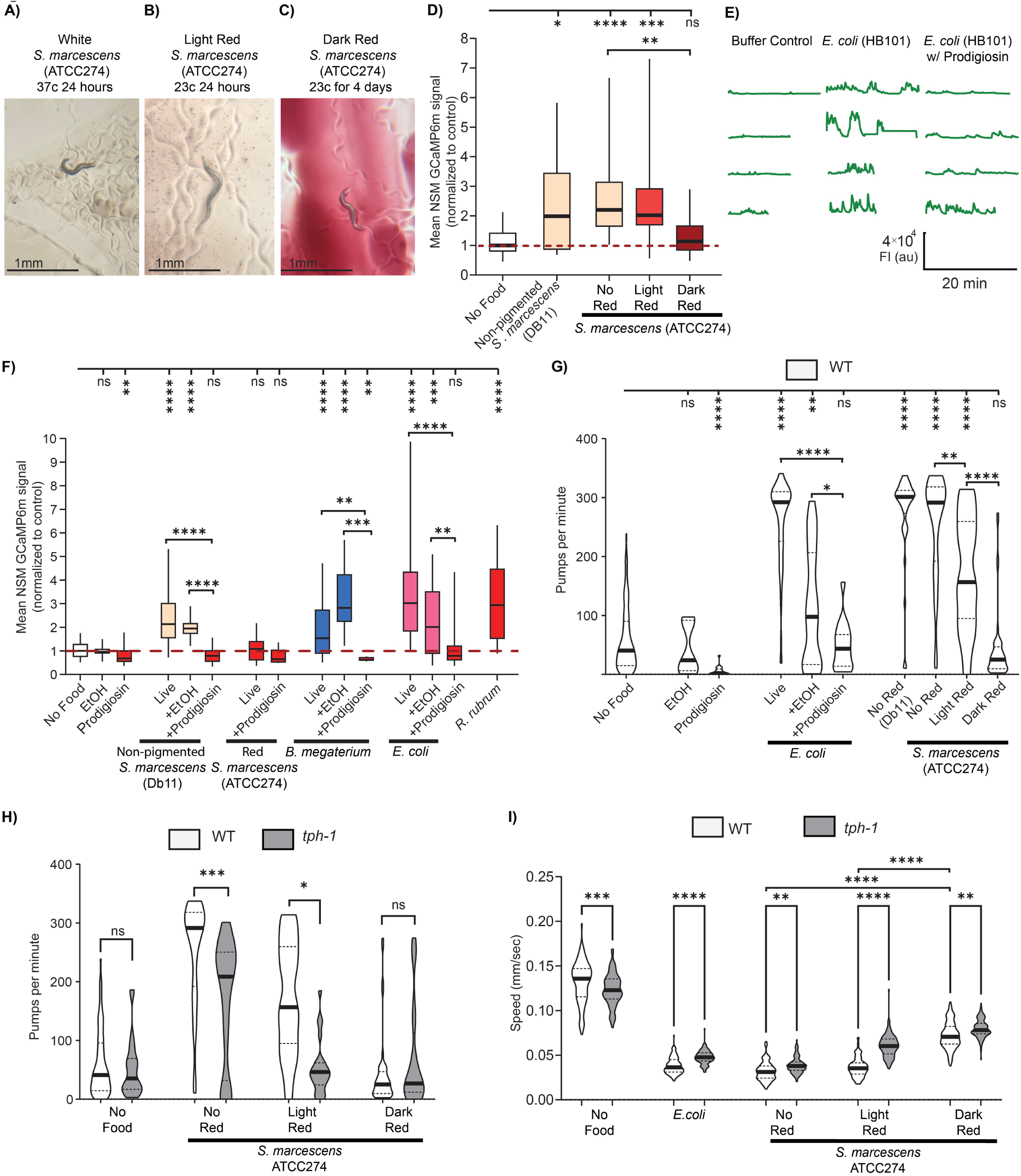
A *Serratia marcescens* metabolite, prodigiosin, inhibits NSM activity and associated behaviors. (A) Representative image of the white color of *S. marcescens* ATCC274 grown at 37C for 24 hours, with a WT animal. (B) Representative image of the light red color of *S. marcescens* ATCC274 grown at 23C for 24 hours, with a WT animal. (C) Representative image of the dark red color of *S. marcescens* ATCC274 grown at 23C for 4 days, with a WT animal. (D) Mean GCaMP signal from NSM in *unc-13(s69)* animals, shown as in Fig. 1C. ****p<0.0001, ***p<0.001, **p<0.005, *p<0.01 by Bonferroni-corrected Mann-Whitney test. n =25-63 animals. (E) Example NSM GCaMP traces from *unc-13(s69)* animals in the indicated conditions. Each line is a recording of one animal (the data are not paired, so different animals are shown for the different experimental conditions). (F) Mean GCaMP signal from NSM in *unc-13(s69)* animals on indicated conditions, shown as in Fig. 1C. Prodigiosin concentration was 200 ug/ml in these experiments, and these results were also replicated at 50 ug/ml and 100 ug/ml in Fig. S3C. ****p<0.0001, ***p<0.001, **p<0.005, *p<0.01 by Bonferroni-corrected Mann-Whitney test. n = 7-170 animals. (G) Pharyngeal pumping rates of WT animals exposed to the indicated substances for one hour. Data displayed as in Fig. 4A. Prodigiosin concentration was 50 ug/ml in these experiments. ****p<0.0001, ***p<0.001, **p<0.005; ns, not significant by Bonferroni-corrected Mann-Whitney test. n = 5-32 animals. (H) Pharyngeal pumping rates of WT (white) and *tph-1* (gray) animals exposed to the indicated substances for one hour, displayed as in Fig. 4A. Note that WT data here is the same data as in (G). ****p<0.0001, ***p<0.001, **p<0.005; ns, not significant by Bonferroni-corrected Mann-Whitney test. n = 10-32 animals. (I) Mean speed for WT (white) and *tph-1*(gray) animals exposed to the indicated substances, displayed as in Fig. 4C. ****p<0.0001; ns, not significant by Mann-Whitney test. n = 100-125 animals per condition across at least three independent days.

We next tested whether the red pigment prodigiosin could potentially explain these effects. Prodigiosin has been shown to affect mammalian cell growth and cell death^67–70^, but its effects on neural circuits have not been examined. In addition, its effects on *C. elegans* behavior have not been investigated. We asked whether the addition of purified prodigiosin (resuspended in ethanol) to appetitive bacteria (*E. coli*, non-pigmented *Serratia*, or *B. megaterium*) could reduce the NSM activation elicited by their ingestion. Indeed, for all three bacterial species (*S. marcescens*, *B. megaterium*, and *E. coli*), the addition of prodigiosin led to a robust suppression of NSM activation (Fig. 5E, F; effect was observed across a range of concentrations, Fig. S3C). This inhibitory effect was not observed for all red pigments, as another red-pigmented bacterium (*R. rubrum*), which produces different bacteriochlorophyll pigments, still robustly activated NSM (Fig. 5F). In addition, this effect was not due to lower pumping rates on prodigiosin, as elevating pharyngeal pumping of animals exposed to *E. coli* plan prodigiosin (via aldicarb addition) did not restore NSM activity (Fig. S3D). Overall, these results suggest that prodigiosin, a metabolite produced by pathogenic *S. marcescens*, inhibits bacterial activation of NSM.

We next examined whether these effects of non-pigmented versus Red *Serratia* could be detected at the behavioral level. To do so, we quantified their effects on pharyngeal pumping and locomotion. Exposure to non-pigmented *Serratia* caused an increase in pharyngeal pumping, but exposure to red *Serratia* did not elevate pharyngeal pumping relative to a no-food control (Fig. 5G). Similarly, addition of prodigiosin to live *E. coli* attenuated the increase in pharyngeal pumping evoked by *E. coli* exposure (Fig. 5G). In addition, exposure to non-pigmented *Serratia* led to a larger reduction in locomotion speed than exposure to Red *Serratia* (Fig. 5I). We next examined whether these behavioral changes required serotonin. *tph-1* mutants had reduced pumping rates relative to wild-type animals when exposed to non-pigmented and light red *Serratia*, indicating a serotonin-dependent response. However, on dark red *Serratia*, where NSM activity appears suppressed, serotonin was no longer necessary for the minimal pumping observed (Fig. 5H). The slow locomotion evoked by these bacteria was also attenuated in *tph-1* mutants (Fig. 5I). Altogether, these results are consistent with our NSM GCaMP imaging experiments above and suggest that *S. marcescens* can have different effects on *C. elegans* behavior depending on its growth conditions and prodigiosin production. This suggests an additional mechanism by which bacteria interact with neurons: by producing metabolites like prodigiosin, bacteria can mask their activating signals and evade neuronal recognition.

## DISCUSSION

Animals couple the microbial contents of their alimentary canals to changes in physiology and behavior, but the signaling mechanisms underlying microbial effects on behavior are not well understood. Here, we found that bacterial polysaccharides in the pharyngeal lumen of *C. elegans* activate NSM through ASIC channels localized to its sensory dendrite. We identify large polysaccharides from *E. coli* and peptidoglycan from gram-positive bacteria as NSM-activating signals. In addition, prodigiosin from infectious *S. marcescens* inhibits bacterial activation of NSM. These results are supported by observing effects on both NSM activity and the feeding and locomotion behaviors that NSM controls. Together, these experiments identify multiple signaling mechanisms that link bacteria of the alimentary canal to neural activity and behavior.

Prior work identified NSM as a neuron that drives appetitive behaviors, like feeding and slow locomotion, in *C. elegans*^36–39^. In addition, prior work had noted its unique morphology and the connection to food intake^36,38,49^. However, the exact features of ingested bacteria that activate NSM were unknown. Our previous work suggested that pharyngeal muscle movements and the flow of small beads through the lumen were insufficient to activate NSM^36^. Here, we found that the ingestion of specific bacterial products (lysates, polysaccharides) is sufficient to activate NSM. This suggests that there is a chemical component to NSM’s detection of bacteria. It is still formally possible that there is a mechanical component as well, since these compounds were actively flowing through the lumen during NSM GCaMP imaging. Prior work has shown that ASICs can be dually sensitive to fluid movement and chemical signals^71^. In addition, a prior study showed that DEL-3 mediates a mechanosensory response in dopaminergic neurons^72^.

We identified peptidoglycan from gram-positive bacteria as one signal that is sufficient to activate NSM. Peptidoglycan is a fairly ubiquitous component of bacterial cell membranes, so NSM’s ability to respond to it may be one of the major reasons that it has such a broad response profile to diverse bacteria. Interestingly, in *Drosophila,* peptidoglycan in the gut has also been shown to modulate host behavior.^74,75^ In addition to peptidoglycan, we also identified prodigiosin from pathogenic *S. marcescens* as an inhibitory signal that can suppress NSM’s response to appetitive bacteria. This suggests an overall model in which NSM responds broadly to polysaccharides present in most bacteria, but also integrates aversive signals that inhibit its activation and modify its overall bacterial response profile. Bacteria that produce masking molecules like prodigiosin may not be recognized by NSM. This ability to evade neuronal recognition may be loosely analogous to bacterial evasion of detection by immune cells.

The activation of NSM by peptidoglycan required the ASICs DEL-3 and DEL-7. However, it remains unclear whether these ASICs are primary sensors of peptidoglycan or, alternatively, whether they are activated by signal transduction after a different primary sensor detects peptidoglycan. In mammals, TLR2, NLRs, and PGLYRPs acts as peptidoglycan sensors. While there is one Toll gene in *C. elegans*, *tol-1*, it does not appear to be expressed in NSM based on transcriptional reporters, single-cell sequencing, or cell-specific mRNA pulldowns^36,76–79^. There are no known NLR or PGLYRP homologs in *C. elegans*^80^. Based on homology, the strongest potential candidates for polysaccharide receptors in NSM are perhaps the C-type lectin receptors (CLRs). In other animals, CLRs play a role in detection of microbial sugars. *C. elegans* has a large family of CLRs (*clec* genes), including some that appear to be expressed in NSM (e.g. *clec-179*)^36,81^. Several *clec* genes are required for pathogen responses in *C. elegans*, but none have been implicated in NSM function^82,83^. Our work here suggests that efforts to identify the full sensory transduction pathway for bacterial polysaccharide detection in NSM is a promising avenue for further study.

In addition to peptidoglycan, there are likely to be other chemical activators of NSM produced by nutritive bacteria. For example, we found that polysaccharides isolated from *E. coli* activated NSM, but we did not identify the exact *E. coli* polysaccharide(s) involved. It is possible that capsular polysaccharides or exopolysaccharides from *E. coli* could act as key signals. Future research will clarify whether NSM responds to select polysaccharide moieties or, alternatively, whether it is more of a promiscuous long-chain polysaccharide sensor.

In their natural habitats, *C. elegans* are thought to eat a wide range of different bacterial species^16^. In this study, we identified microbial signals that activate NSM, namely polysaccharides, and at least one that inhibits NSM activation, prodigiosin from infectious *S. marcescens*. Given that the animal might often be ingesting a mixture of microbial signals, NSM may act as a hub neuron that integrates information about the microbial contents of its pharyngeal lumen. Its release of serotonin can influence activity in a large fraction of the neurons of the *C. elegans* nervous system^60^, allowing it to broadcast information about the contents of lumen to the rest of the brain.

## ACKNOWLEDGMENTS

We thank Piali Sengupta, Michael O’Donnell, Likui Feng, Cori Bargmann, and members of the Flavell lab for critical reading of the manuscript. We thank Gary Ruvkun and the CGC (supported by P40 OD010440) for sharing bacterial and *C. elegans* strains. C.E. acknowledges funding from NIH (F32 NS116107) and a JPB Foundation Postdoctoral Fellowship. S.W.F. acknowledges funding from NIH (GM135413, DC020484); the McKnight Foundation; Alfred P. Sloan Foundation; The Picower Institute for Learning and Memory; the Howard Hughes Medical Institute; and The Freedom Together Foundation. S.W.F. is an investigator of the Howard Hughes Medical Institute.

## AUTHOR CONTRIBUTIONS

Conceptualization, C.E. and S.W.F. Methodology, C.E., M.D., C.P.F., G.J.H., M.A., B.L.G, L.L.D. Software, C.E. and C.P.F. Formal analysis, C.E. Investigation, C.E., M.D., M.A., L.L.D. Writing – Original Draft, C.E. and S.W.F. Writing – Review & Editing, C.E., C.P.F., G.J.H., B.L.G., L.L.D. and S.W.F. Funding Acquisition, B.L.G. and S.W.F.

## MATERIALS AND METHODS

### Key Resources Table

| REAGENT or RESOURCE | SOURCE | IDENTIFIER |
| --- | --- | --- |
| Experimental Models: Organisms/Strains |  |  |
| *C. elegans*: Wild-type Bristol N2 | Caenorhabditis Genetics Center | N2 |
| *C. elegans*: *unc-13(s69); flvls146[Pdel-7::GCaMP6m, myo2::mCherry]* | This study | SWF856 |
| *C. elegans*: *del-7(ok1187); del-3(ok2613); unc-13(ky1027); lite-1(ce314); flvls146[Pdel-7::GCaMP6m, myo2::mCherry]* | This study | SWF814 |
| *C. elegans*: *tph-1 (mg280) II* | Caenorhabditis Genetics Center | MT15434 |
| Bacterial and Virus Strains |  |  |
| *Escherichia coli*: Strain OP50 | Caenorhabditis Genetics Center | OP50 |
| *Escherichia coli* Strain HB101 | Caenorhabditis Genetics Center | HB101 |
| *Delftia acidovorans*: Strain JUb08 | Samuel et al, 2016 | JUb08 |
| *Bacillus subtilis*: Strain JUb10 | Samuel et al, 2016 | JUb10 |
| *Comamonas sp.*: Strain DA1877 | Caenorhabditis Genetics Center | DA1877 |
| *Staphylococcus aureus*: Strain JUb015 | Samuel et al, 2016 | JUb015 |
| *Stenotrophomonas maltophilia*: Strain JUb023 | Samuel et al, 2016 | JUb023 |
| *Achromobacter denitrificans*: Strain JUb024 | Samuel et al, 2016 | JUb024 |
| *Sphingobacterium*: Strain JUb094 | Samuel et al, 2016 | JUb094 |
| *Pseudomonas aeruginosa*: Strain JUb095 | Samuel et al, 2016 | JUb095 |
| *Lactococcus*: Strain JUb097 | Samuel et al, 2016 | JUb097 |
| *Leucobacter*: Strain BIGb0196 | Samuel et al, 2016 | BIGb0196 |
| *Chryseobacterium indologenes*: Strain BIGb0234 | Samuel et al, 2016 | BIGb0234 |
| *Serratia marcescens*: Strain ATCC274 | ATCC | ATCC274 |
| *Microbacterium nematophilum*: Strain CBX102 | Caenorhabditis Genetics Center | CBX102 |
| *Serratia marcescens*: Strain Db11 | Caenorhabditis Genetics Center | Db11 |
| *Bacillus megaterium*: Strain DA1880 | Caenorhabditis Genetics Center | DA1880 |
| *Bacillus cereus* | Home Science Tools | LD-BACGRAM |
| *Branhamella catarrhalis* | Home Science Tools | LD-BACGRAM |
| *Micrococcus luteus* | Home Science Tools | LD-BACGRAM |
| *Rhodospirillum rubrum* | Home Science Tools | LD-BACGRAM |
| *Serratia marcescens* | Home Science Tools | LD-BACGRAM |
| Chemicals, Peptides, and proteins |  |  |
| Bovine Serum Albumin | Sigma | A8806 |
| Fisher BioReagents™ Microbiology Media Additives: Peptone | Fisher scientific | BP1420 |
| Yeast Synthetic Drop-out Medium Supplements | Millipore Sigma | Y2001 |
| Peptidoglycan from *Bacillus subtilis* | Millipore Sigma | 69554 |
| PGN-ECndss ultrapure, soluble | Invivogen | tlrl-ksspgn |
| Peptidoglycan from *Micrococcus luteus* | Millipore Sigma | 53243 |
| *E. coli* Extract Total | Avanti Research | 100500P-100mg |
| Ultra Omega-3++ from Fish Oil 1400 mg Softgels | Nature Made | B0046XC528 |
| D-(+)-Glucose | Millipore Sigma | G8270 |
| D-(+)-Xylose | Millipore Sigma | X1500-10MG |
| D-(+)-Maltose monohydrate | Millipore Sigma | M5885-100G |
| L-(+)-Arabinose | Millipore Sigma | A3256-10MG |
| D-Glucuronic acid | Millipore Sigma | G5269-10MG |
| D-(+)-Galactose | Millipore Sigma | G0625-100G |
| D-(–)-Fructose | Millipore Sigma | F0127-10MG |
| Sucrose | Millipore Sigma | S8501 |
| D-(+)-Raffinose pentahydrate | Millipore Sigma | R0250-25G |
| Glucose, Diglycerides | Coffee-Mate Original | B00MB2WJTU |
| N-Acetylneuraminic acid | Millipore Sigma | A2388-10MG |
| N-Acetyl-D-mannosamine | Millipore Sigma | A8176-10MG |
| N-Acetyl-D-glucosamine | Mpbio | 7512-17-6 |
| 2-Keto-3-deoxyoctonate ammonium salt | Millipore Sigma | K2755 |
| Glycogen, from Oysters, Ultrapure, Thermo Scientific | Fisher Scientific | AAJ1644506 |
| Mannan from Saccharomyces cerevisiae | Millipore Sigma | M7504-100MG |
| Lipopolysaccharides from Escherichia coli O55:B5 | Millipore Sigma | L3012 |
| Lipopolysaccharides from Escherichia coli O26:B6 | Millipore Sigma | L2762 |
| Pro-Q™ Emerald 300 Glycoprotein Gel and Blot Stain Kit | Thermo Fisher | P21857 |
| FluoSpheres™ Carboxylate-Modified Microspheres, 0.02 µm, red fluorescent (580/605), 2% solids | Thermo Fisher | F8786 |
| Software and Algorithms: |  |  |
| Adobe Illustrator | Adobe | https://www.adobe.com |
| Matlab (R2020b, R2021a, R2014a) | Mathworks | www.mathworks.com |
| Prism (v10) | Graphpad | www.graphpad.com |
| ImageJ (v1.50) | NIH | http://imagej.nih.gov/ij/ |
| NIS-Elements (v4.51.01) | Nikon | www.nikoninstruments.com/products/software |
| Streampix (v7.0) | Norpix | https://www.norpix.com |
| Prime BSI sCMOS camera | Photometrics | N/A |
| Zyla 4.2 Plus sCMOS camera | Andor | N/A |
| X-Cite 120LED System XT120L | Lumen Dynamics | N/A |
| Ti-S Inverted Microscope | Nikon | N/A |
| SP-20000M-USB3 CMOS camera | JAI | N/A |
| 10x25 White Panel LED backlight, 24VDC | Metaphase Technologies | Cat#MS-BL10X25-W-24-ILD-PS |
| ZEISS Stemi 305 | Zeiss | N/A |

### *C. elegans* growth and genetics

*C. elegans* Bristol strain N2 was used as wild-type. All wild-type, mutant and transgenic strains used are listed in the key resources table. Animals were raised on NGM agar plates seeded with *E. coli* OP50 bacteria and kept at room temperature (∼23°C). Young adults were used for all experiments. For genetic crosses, genotypes were confirmed by PCR and/or sequencing. Transgenic animals were generated by injecting DNA with fluorescent co-injection markers into the gonads of young adult hermaphrodites. Transgene integration was by UV/TMP and the integrated strain was backcrossed 6 times after integration.

### Calcium Imaging

Calcium imaging was conducted as previously described^36^ with minor modifications. Experimental substances (bacteria, macromolecules, etc) were pipetted (6 µL) onto flat NGM agar pads mounted on glass slides and allowed to dry before the animals were introduced. Animals were removed from food for at least 15 minutes prior to the assay. After transferring animals onto the dried substance, polydimethylsiloxane (PDMS) arenas were positioned around each animal and the preparations were enclosed with cover glass. GCaMP6m fluorescence was recorded at 10 fps using a 4×/0.2 NA objective and an Andor Zyla 4.2 Plus sCMOS cameras (or Photometrics Prime BSI sCMOS, for some recordings). For analysis, the NSM soma was identified using the GCaMP6m signal and tracked using custom MATLAB scripts. Fluorescent signals from a region-of-interest (ROI) containing the soma were extracted at each time point. Background fluorescence was subtracted, and the mean GCaMP6m intensity within each ROI was used as the data point for each animal. As is described in the figure legends, data were normalized to day-matched no-food control recordings.

Live bacteria were tested at various optical densities (OD₆₀₀) ranging from 0.01 to 3.0. Specifically, the ODs tested were 0.01, 0.05, 0.1, 0.5, 0.6, 1.0, 2.0, and 3.0. Because no significant difference in NSM activity was observed at ODs above 0.5, these concentrations were pooled for analysis. An amino acid solution in LB was tested at concentrations of 1 mg/mL and 20 mg/mL. Peptone was tested at 2, 100, and 500 mg/mL. Bovine serum albumin (BSA) was tested at 125, 250, 1000, and 2000 µg/mL. *E. coli* lipids, both commercially obtained from Avanti Polar Lipids and lab-purified, were tested at 0.1, 1, and 10 mg/mL. Fish oil was tested at 0.5, 2, and 600 mg/mL. Lipopolysaccharides (LPS) from *E. coli* strains O126:B8 and O55:B5 were tested at 20 µM and 10 mM. Glucose was tested at 20, 100, 250, and 500 mM. Glucose combined with diglycerides was tested at 5, 500, and 1000 mg/mL. Purified *E. coli* DNA was tested at 400 ng/µL. Glucose, galactose, mannose, maltose, arabinose, xylose, sucrose and raffinose were tested at 20, 250, 500 mM, and 1M. Gluconic acid and sialic acid were tested at 500mM. NAM and NAG were tested at 30, 200 and 800mM. Sorbitol tested at 50 and 250 mM. KDO tested at 2 and 4 mM. Glycogen tested at 100mg/ml and mannan at 2mg/ml. For each of these compounds, each concentration was tested separately and then concentrations were pooled in the presented graphs after no significant difference between concentrations was found using a non-parametric Mann-Whitney test.

For acid hydrolysis treatment of clarified lysate and purified polysaccharides, the samples were treated with 3N Hydrochloric acid until they reached a pH of 1, and were then incubated at room temperature for at least 3 hours. Then 10M sodium hydroxide was added until the solution reached a pH of 7. This sample was then tested for NSM activity.

### Behavior Experiments

All behavioral experiments (speed and feeding assays) were done over at least two separate days and over multiple experimental replicates. All animals were staged as L4s the day before the experiment.

### Animal Speed Recordings

Multi-worm tracking was performed as previously described^36^. One-day-old adult animals of the indicated genotypes were transferred to 10 cm² round NGM agar plates that had been seeded with 250 µL of the experimental substance (bacteria, macromolecule, etc). When the experimental substance contained live bacteria, the optical density (OD₆₀₀) was adjusted to 2.0 prior to application on the agar surface. Plates were allowed to dry completely before animals were introduced. For behavioral assays monitoring locomotion, animals were fasted for at least 30 minutes by transferring them to unseeded NGM plates (lacking *E. coli* OP50) prior to recording and the recordings were 60 minutes long. Animal speed was averaged across time for each isolated worm track to determine each animal’s average speed in a given environmental condition. Videos were captured using Streampix software at 3 frames per second (fps) with JAI SP-20000M-USB3 CMOS cameras (5120 × 3840, monochrome) equipped with Nikon Micro-NIKKOR 55 mm f/2.8 lenses. White-panel LEDs (Metaphase) provided backlighting. Tracking data were analyzed using custom MATLAB scripts as previously described^36^.

### Feeding assays

Pumping assays were performed on NGM plates containing varying concentrations of experimental substances. For wild-type and *tph-1* mutant pumping assays, plates were seeded with 250 µL of the test solution to cover the entire agar surface and plates were then allowed to dry. Young adults (1–10 per plate) were transferred onto plates, allowed to acclimate for 30-60 minutes, and then pumping was counted under a Zeiss Stemi 305 microscope. The free app Tap Tool v.1.3 was used to tally pumps.

### Aldicarb Treatment for NSM Imaging in *unc-13* Mutants

To perturb pharyngeal pumping behavior during NSM calcium imaging, aldicarb—a well-characterized acetylcholinesterase inhibitor—was applied to the surface of imaging plates. Although cholinergic transmission is significantly reduced in *unc-13* mutants, application of 0.5 mM aldicarb was sufficient to significantly increase pumping, while 26 mM aldicarb led to a robust inhibition of pumping (Fig. S1A–B). For each assay, 6 µL of the aldicarb-containing solution was added onto the agar surface of the imaging slide and allowed to dry completely. The next solution was added on top of the dried spot, for example 6 µL of *E. coli* or prodigiosin. After the liquid had dried, the young adult *unc-13* mutant animals were transferred to the treated agar and allowed to acclimate for 5 minutes. Pumping rates were quantified visually under a Zeiss Stemi 305 dissection microscope using the free app Tap Tool v.1.3. After counting pumps, animals were subject to NSM calcium imaging as described above.

### Bacterial strain and growth conditions

All bacterial strains were streaked to single colonies on LB plates. A single colony was inoculated in 4 ml of LB in a 15 ml round bottom tube with loose cap to allow for adequate aeration and grown at 37 °C under shaking conditions for 16 hours. All bacteria were stored at 4°C. Prior to application on NGM plates, the OD was adjusted as described above.

### Prodigiosin

Prodigiosin was dissolved in ethanol at a concentration of 200 ug/ml. In experiments with prodigiosin, bacteria at an OD₆₀₀ = 1 was added to the NGM agar surface and allowed to dry. Then, an equal amount of prodigiosin at a concentration of 50-200 ug/ml (as indicated in figure legends) was added on top of the bacteria spot. The negative control for prodigiosin was ethanol alone (without prodigiosin added).

### Fluorescent Bead Ingestion

To test whether animals were ingesting the contents of the agar surface, we added fluorescent nanobeads (Red Fluorospheres with bead size 0.02um, Invitrogen) to the agar surface. After allowing animals (strain: *unc-13(s69); flvIs146[Pdel-7::GCaMP6m, myo2::mCherry]*) to ingest beads for 10 minutes, we imaged the animals under a TRITC filter to determine the red fluorescent bead location.

### GC–MS Analyses of Derivatized Sugars

Gas chromatography-mass spectrometry (GC-MS) analysis was performed using an Agilent 7890A GC, coupled with a 5975C MS detector (MSD) capable of scanning up to 1000 m/z. The system utilized electron impact (EI) ionization at an approximate voltage of 1900 V, with data acquisition and analysis conducted using ChemStation software (Agilent Technologies). Prior to analysis, bacterial carbohydrate samples underwent trifluoroacetic acid (TFA) hydrolysis (2M) for varying durations of 1.5, 3, and 5 hours to release monosaccharides. Following hydrolysis, samples were derivatized using trimethylsilyl (TMS) reagents to enhance volatility for GC-MS analysis. Derivatization was carried out by adding 1-(Trimethylsilyl)-1 H-imidazole (TMSI) Silylation Reagent (ThermoFisher Scientific), followed by incubation at 80°C for 30–60 minutes. After cooling to room temperature, the samples were injected into the GC-MS system at 270°C using a 20:1 split mode for analysis.

For gas chromatographic separation, a DB-5HT MS column (30 m × 0.25 mm ID × 0.25 µm film thickness with a maximum temperature limit of 400°C) was used with helium (99.999%) as the carrier gas at a flow rate of 0.6 mL/min. The oven temperature was programmed to increase from 40°C to 375°C, with the gradient optimized for effective separation. The mass spectrometer was operated with a scan range of 45–900 m/z. The ion source temperature was set to 230°C, and the quadrupole temperature was maintained at 150°C. Peaks were identified by comparing retention times to sugar standards and by matching the mass profiles of the sugar fragments against the NIST and Wiley spectral libraries, allowing for compound confirmation based on mass fragmentation patterns. Data analysis involved integrating peak areas and compiling them into a summary table, which was subsequently used to calculate the relative ratios of detected sugars. The raw mass spectrometry data is available at: https://www.dropbox.com/scl/fo/n6t4or1nv38lhs6dnaiuo/AIkj_4zm7TIzkEAwOfmk9sw?rlkey=lohg2c4vieykjgjhw6zwekkuh&dl=0

### *E. coli* Polysaccharide Purification

For biochemistry experiments, we used *E. coli* strain HB101 [supE44 hsdS20(rB-mB-) recA13 ara-14 proA2 lacY1 galK2 rpsL20 xyl-5 mtl-1] and used the following protocol. HB101 was grown in 1 liter of Terrific Broth and shaken at 37°C overnight. The next day the bacteria was spun down at 8,000 x g for 15 minutes. The pellet was washed four times with distilled water and resuspended in 50 ml ice cold PBS along with 100ul DTT (1M), 100ul Aprotinin (100x), 30ul Dnase1 (10mg/ml). The HB101 solution was then put through the LM20 high-pressure microfluidizer-3 passes at 20000 PSI then 3 passes at 25000K PSI. Next, the HB101 homogenized cell lysate was put through a high-speed spin using a 45Ti rotor in an ultracentrifuge at 150,000-180,000 x g for 2 hours. After centrifugation both the supernatant and pellet were collected and stored at 4°C. From the high-speed supernatant (now labeled ‘clarified lysate’), we performed an ethanol precipitation at -20°C overnight. The final volume of ethanol was either 5% or 70%, as indicated in figure legends. The ethanol solution was spun in a microcentrifuge at 13.3 rpm for 10 minutes. After centrifugation of ethanol-treated solution, precipitates were collected and resuspended in 1x PBS at pH 7.4 at the same original volume of the high-speed spin supernatant used. To digest proteins, 800 units/ml of proteinase K (a broad-spectrum serine protease) was added to the resuspended ethanol precipitate for 1 hour at 37°C. Protein digestion was confirmed by running samples on Any kD Mini-PROTEAN TGX Stain-Free protein gels. For DNA digestion, 10 units/ml of DNase 1, was added to the resuspended ethanol precipitate for 1 hour at 37°C. For RNA digestion, 1 units/ml of RNase H was added to the resuspended ethanol precipitate for 1 hour at 37°C.

### Statistics

Non-parametric statistical tests were used throughout the study. Bonferroni correction was applied to correct for multiple comparisons when using pairwise tests. For some large group analyses, we used FDR correction, as indicated in the figure legends.

---

**Supplementary Figure 1.**
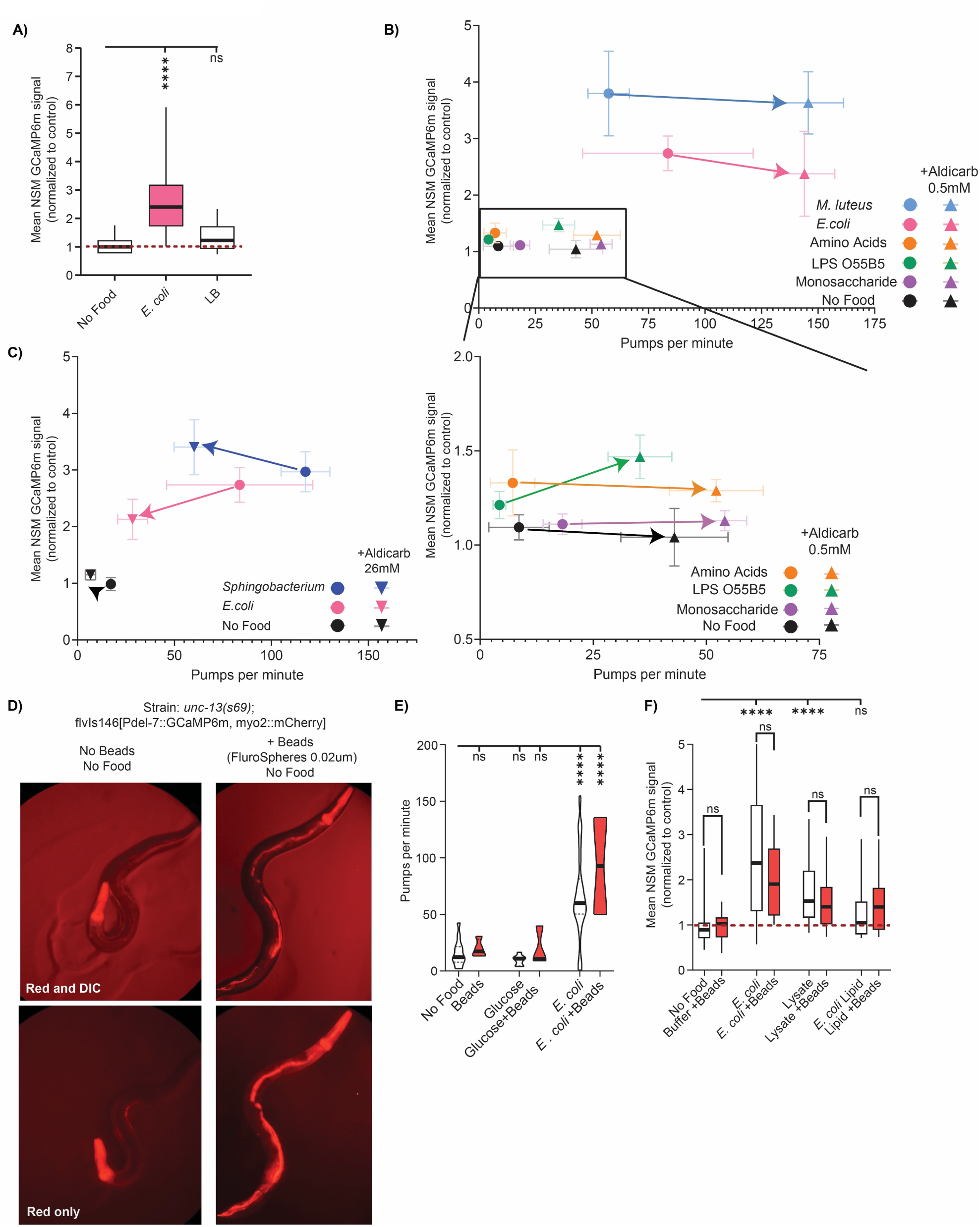
(A) Mean GCaMP signal from NSM in *unc-13(s69)* animals, including Luria-Bertani (LB) broth negative control, shown as in Fig. 1C. ****p<0.0001 by Bonferroni-corrected Mann-Whitney test. n =12-45 animals. (B) Scatter plot showing relationship between pumping and NSM activity (see main text for rationale). Circle dots represent mean NSM GCaMP signals from *unc-13(s69)* animals under various conditions without aldicarb. Triangles represent mean NSM GCaMP signals from animals treated with 0.5 mM aldicarb to increase pumping. Arrows connect data from the same experimental conditions where the only difference is aldicarb addition. Top: zoomed out plot. Bottom: a zoomed-in view of the inset box for clarity. Error bars are SEM. n=7-20 animals. (C) Scatter plot showing relationship between pumping and NSM activity (see main text for rationale). Circle dots represent mean NSM GCaMP signals from *unc-13(s69)* animals under various conditions without aldicarb. Inverted triangles represent NSM signals from animals treated with 26 mM Aldicarb to reduce pumping. Arrows connect data from the same experimental conditions where the only difference is aldicarb addition. Error bars are SEM. n=11-22 animals. (D) Representative images of *unc-13(s69)*; *flvIs146*[*del-7::GCaMP6m, myo2::mCherry*] animals with and without addition of FluroSphere 0.02um red fluorescent beads to the agar surface. These images were collected in the absence of food, in order to test whether the low basal pumping rate in the absence of food is sufficient for animals to ingest the contents on the agar surface. The +Beads images (right) show that red beads are clearly visible in the intestinal tract of the animal even under these no food conditions, indicating that substantial ingestion occurs even with a fairly low pumping rate. Note that the red color in the pharynx is the fluorescent co-injection marker also present in the absence of beads (left). Top images include the red and DIC channel to show the body outline of the animals; bottom images just show red fluorescence. Animals were permitted to ingest beads for 10 minutes prior to imaging (see Methods) (E) Pharyngeal pumping rates of *unc-13(s69)* animals, exposed to the indicated substances for 15 minutes. n = 8-45 animals. (F) Mean GCaMP signal from NSM in *unc-13(s69)* animals, for the indicated experimental conditions, which include the addition of the beads shown in (D). n =8-30 animals.

**Supplementary Figure 2.**
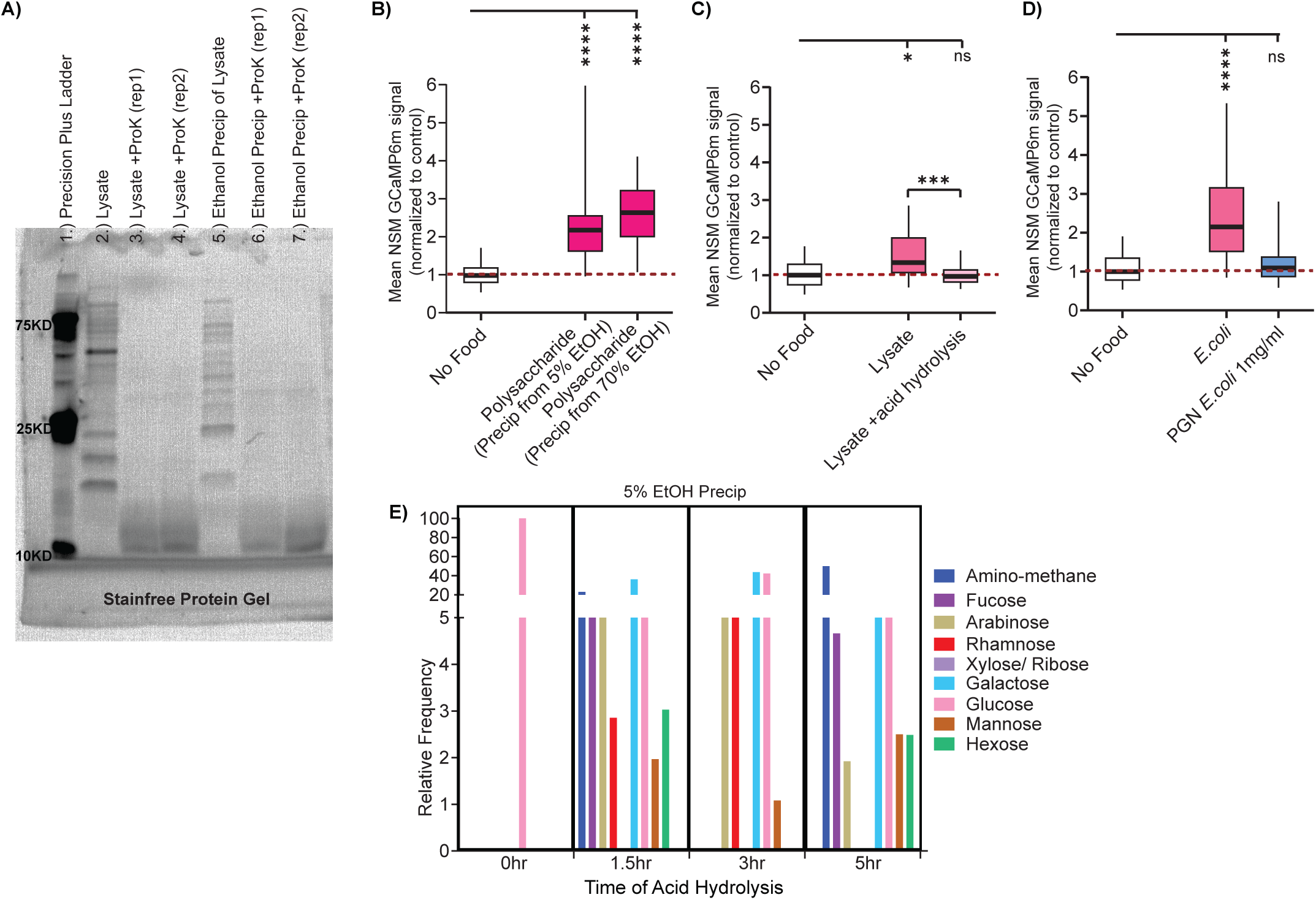
(A) Stainfree protein gel showing proteinase K digestion of clarified lysate (compare lane 2 with lanes 3 and 4) and ethanol precipitated sample subjected to RNase, DNase, and Proteinase K treatment (compare lane 5 with lanes 6 and 7). (B) Mean GCaMP signal from NSM in *unc-13(s69)* animals on indicated conditions, shown as in Fig. 1C. ****p<0.0001 by Mann-Whitney test. n = 35-48 animals. (C) Mean GCaMP signal from NSM in *unc-13(s69)* animals on indicated conditions, shown as in Fig. 1C. ***p<0.001, *p<0.01 by Mann-Whitney test. n = 38-45 animals. (D) Mean GCaMP signal from NSM in *unc-13(s69)* animals on indicated conditions, shown as in Fig. 1C. ****p<0.0001 by Mann-Whitney test. n = 42-120 animals. (E) GC-MS carbohydrate profiling of a purified bacterial polysaccharide sample, shown as shown in Fig. 2D. Whereas the dataset in Fig. 2D was from a 70% ethanol precipitation, this sample was precipitated with 5% ethanol; the results are qualitatively similar across both samples.

**Supplementary Figure 3.**
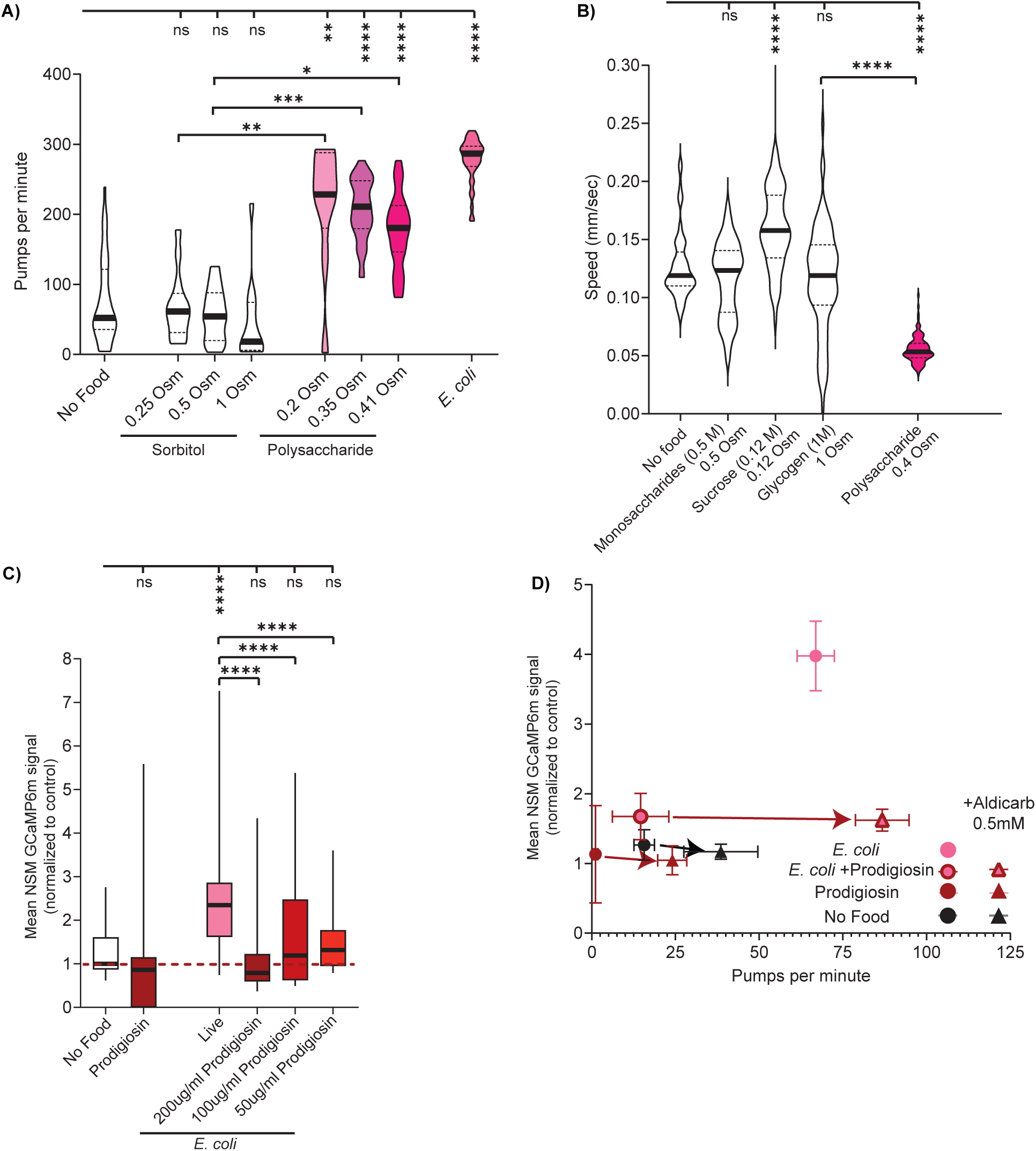
(A) Pharyngeal pumping rates of WT animals exposed to the indicated substances for one hour, shown as in Fig. 4A. ****p<0.0001, ***p<0.001, *p<0.01; ns, not significant by Bonferroni-corrected Mann-Whitney test. n = 9-86 animals. (B) Mean speed for WT animals on indicated substances, shown as in Fig. 4C. Monosaccharides condition combines separate experiments of 0.5 M Glucose, Galactose, and Maltose. ****p<0.0001, ***p<0.001, **p<0.005, *p<0.01, by Bonferroni-corrected Mann-Whitney test. N = 15-125 animals per condition across at least three independent days. (C) Mean GCaMP signal from NSM in *unc-13(s69)* animals on indicated conditions, shown as in Fig. 1C. ****p<0.0001 by Mann-Whitney test. n = 10-51 animals. (D) Scatter plot showing that increased pumping does not increase NSM activity in the presence of *E. coli* + prodigiosin. Circle dots represent mean NSM GCaMP signals from *unc-13(s69)* animals under various conditions without aldicarb. Triangles represent mean NSM GCaMP signals from animals treated with 0.5 mM aldicarb to increase pumping. Arrows connect data from the same experimental conditions where the only difference is aldicarb addition. Error bars are SEM. n=8-44 animals.

**Supplementary Table 1.** Table 1 shows the characteristics for all the bacteria used in this study. Activity in NSM column is the average fluorescent intensity fold increase over the negative control (i.e. No Food). Strain class column is from Samuel et al., unless otherwise referenced.

| Bacteria | Activity in NSM (x over No Food) | Size (µm) | Morphology | Pathogenicity | Gram-Stain | Phylum | Class | Order | Family |
| --- | --- | --- | --- | --- | --- | --- | --- | --- | --- |
| *E. coli* | 2x | 1 x 2-4 | Rod (Bacillus) | Opportunistic | Gram-negative | Proteobacteria | Gammaproteobacteria | Enterobacterales | Enterobacteriaceae |
| *Rhodospirillum rubrum* | 2.7x | 0.8-1.0 x 2-5 | Spiral (Spirillum) | Non-pathogenic | Gram-negative | Proteobacteria | Alphaproteobacteria | Rhodospirillales | Rhodospirillaceae |
| *Sphingobacterium spp.* | 2.4x | 0.5-1 x 2-5 | Rod (Bacillus) | Opportunistic | Gram-negative | Bacteroidota | Sphingobacteriia | Sphingobacteriales | Sphingobacteriaceae |
| *Stenotrophomonas spp.* | 2.3x | 0.7-1.3 x 1.5-3 | Rod (Bacillus) | Opportunistic | Gram-negative | Proteobacteria | Gammaproteobacteria | Xanthomonadales | Xanthomonadaceae |
| *Chryseobacterium spp.* | 2.3x | 0.5-1 x 1-4 | Rod (Bacillus) | Opportunistic | Gram-negative | Bacteroidota | Flavobacteriia | Flavobacteriales | Weeksellaceae |
| *Serratia marcescens* | 2x | 0.5-1 x 0.9-2 | Rod (Bacillus) | Opportunistic | Gram-negative | Proteobacteria | Gammaproteobacteria | Enterobacterales | Enterobacteriaceae |
| *Pseudomonas aeruginosa* | 2x | 0.5-0.8 x 1.5-3 | Rod (Bacillus) | Opportunistic | Gram-negative | Proteobacteria | Gammaproteobacteria | Pseudomonadales | Pseudomonadaceae |
| *Delftia* | 1.6x | 0.5-1 x 2-5 | Rod (Bacillus) | Opportunistic | Gram-negative | Proteobacteria | Betaproteobacteria | Burkholderiales | Comamonadaceae |
| *Achromobacter spp.* | 1.6x | 0.5-1 x 1-3 | Rod (Bacillus) | Opportunistic | Gram-negative | Proteobacteria | Betaproteobacteria | Burkholderiales | Alcaligenaceae |
| *Comamonas aquatica* | 1.4x | 0.3-0.8 x 1.5-3 | Rod (Bacillus) | Non-pathogenic | Gram-negative | Proteobacteria | Betaproteobacteria | Burkholderiales | Comamonadaceae |
| *Staphylococcus spp.* | 2.5x | 0.5-1 | Spherical (Coccus) | Opportunistic | Gram-positive | Firmicutes | Bacilli | Bacillales | Staphylococcaceae |
| *Leucobacter spp.* | 2.5x | 0.5-1 x 1-3 | Rod (Bacillus) | Non-pathogenic | Gram-positive | Actinobacteria | Actinobacteria | Micrococcales | Microbacteriaceae |
| *Lactococcus spp.* | 2.4x | 0.5-1.2 | Spherical (Coccus) | Non-pathogenic | Gram-positive | Firmicutes | Bacilli | Lactobacillales | Streptococcaceae |
| *Micrococcus luteus* | 2.2x | 0.5-3 | Spherical (Coccus) | Non-pathogenic | Gram-positive | Actinobacteria | Actinobacteria | Micrococcales | Micrococcaceae |
| *Microbacterium nematophilum* | 2x | 0.5-1 x 1-3 | Rod (Bacillus) | Non-pathogenic | Gram-positive | Actinobacteria | Actinobacteria | Micrococcales | Microbacteriaceae |
| *Bacillus megaterium* | 2x | 1.5-4 x 5-20 | Rod (Bacillus) | Non-pathogenic | Gram-positive | Firmicutes | Bacilli | Bacillales | Bacillaceae |
| *Bacillus cereus* | 2x | 1 x 3-4 | Rod (Bacillus) | Pathogenic (foodborne) | Gram-positive | Firmicutes | Bacilli | Bacillales | Bacillaceae |
| *Bacillus subtilis* | 2x | 0.7-0.8 x 2-3 | Rod (Bacillus) | Non-pathogenic | Gram-positive | Firmicutes | Bacilli | Bacillales | Bacillaceae |

